# Sharp cell-type-identity changes differentiate the retrosplenial cortex from the neocortex

**DOI:** 10.1101/2022.07.04.498750

**Authors:** Kaitlin E. Sullivan, Larissa Kraus, Lihua Wang, Tara R. Stach, Andrew L. Lemire, Jody Clements, Mark S. Cembrowski

**Affiliations:** Dept. of Cellular and Physiological Sciences, Life Sciences Institute, University of British Columbia, 2350 Health Sciences Boulevard, Vancouver, BC, Canada; Djavad Mowafaghian Centre for Brain Health, University of British Columbia, 2215 Wesbrook Mall, Vancouver, BC, Canada; School of Biomedical Engineering, Biomedical Research Centre, University of British Columbia, 2222 Health Sciences Mall, Vancouver, BC, Canada; Janelia Research Campus, HHMI, 19700 Helix Dr, Ashburn, VA, USA

## Abstract

The laminae of the neocortex are fundamental processing layers of the mammalian brain. Notably, such laminae are believed to be relatively stereotyped across short spatial scales, such that shared laminae between nearby brain regions exhibit similar constituent cells. Here, we considered a potential exception to this rule by studying the retrosplenial cortex (RSC), a brain region known for sharp cytoarchitectonic differences across its granular-dysgranular border. Using a variety of transcriptomics techniques, we identified, spatially mapped, and interpreted the excitatory cell-type landscape of the mouse RSC. In doing so, we surprisingly uncovered that RSC gene expression and cell types change sharply at the granular-dysgranular border. Additionally, supposedly homologous laminae between the RSC and neocortex are effectively wholly distinct in their cell-type composition. In collection, the RSC exhibits a variety of intrinsic cell-type specializations, and embodies a new neocortical organizational principle wherein cell-type identities vary sharply within and between brain regions.

## INTRODUCTION

The neocortex is a recently evolved structure of the mammalian nervous system, with different regions of the neocortex involved in sensory representations, perception, reasoning, motor commands, and many other cognitive abilities^1^. In contrast to this functional diversity, neocortical structure is characterized by a relatively simple 6-layer architecture. Of note, this architecture is believed to be relatively conserved across cortical brain regions^2^, with changes in laminar properties typically occurring in a spatially gradual manner across multiple regions^3,4^. Thus, to a first approximation, neighbouring brain regions of the cortex are thought to share relatively invariant cell-type identities, and thus likely also produce similar intrinsic computations derived from these cell types.

The retrosplenial cortex (RSC), a dorsal midline brain region often studied within a neocortical context^5-7^, may be an exception to these stereotyped cell-type-identity rules. The RSC is commonly split into two broad subregions known as the dysgranular RSC (dRSC) and the granular RSC (gRSC), which show differences between their respective cytoarchitecture. In particular, the 5-layered dRSC is defined by the lack of a layer 4^8^, whereas the adjacent 6-layered gRSC exhibits a clear layer 4 and a relatively dense layer 2/3. Such striking layered differences between the dRSC and gRSC may be suggestive that the shared and supposedly homologous layers between the dRSC and gRSC, and potentially the neocortex, may also vary in their respective cell-type composition.

Revealing the extent and rules of this potential cell-type heterogeneity is important, as this will provide a key conceptual consideration linking cortical organization to function. Of note, the RSC has been hypothesized to assist in switching between egocentric and allocentric frames of reference^8,9^, with the dRSC and gRSC differentially involved in these allocentric and egocentric components. Similar dRSC-gRSC functional dissociations have been observed for a variety of other behavioural settings^6,10,11^. Thus, the dRSC and gRSC subregions demonstrate distinct functional niches, which in principle may be driven by underlying differences in cell-type identity.

As both cytoarchitecture and functional properties change across the RSC, we were motivated to investigate whether such differences might be reflected in specialized rules governing RSC cell-type identity. Using a complement of transcriptomics techniques, we identified numerous genes with RSC-specific and RSC-subregion-specific expression. Expression of these genes delineates spatially restricted excitatory cell types within the RSC, with multiple laminae of the RSC having specialized intrinsic identities that vary dramatically from neocortical counterparts. These findings illustrate that the RSC differs from the cell-type-specific composition and organizational rules of the neocortex, and introduce the principle that supposedly shared laminae across cortical brain regions can actually vary sharply in cell-type identify.

## RESULTS

### The cell-type organization of the RSC via single-cell RNA sequencing

To understand the cell-type landscape of the retrosplenial cortex (RSC), we began by performing single-cell RNA sequencing. We microdissected the anterior, intermediate, and posterior RSC from coronal sections of three animals (Figure S1A). Microdissected tissue was dissociated, and individual cells with neuron-like cell bodies were manually captured. Captured cells underwent library preparation, sequencing, and analysis according to previous methods^12^. From the obtained single-cell transcriptomes, we screened for excitatory neurons with high read counts (see Methods), ultimately resulting in scRNA-seq data from 765 total cells (5.4 ± 1.0 thousand expressed genes/cell from 138 ± 84 thousand reads/cell, mean ± SD; Figure 1A). These data are available online, along with a user interface containing analysis and visualization tools (http://scrnaseq.janelia.org/rsc).

**Figure 1.**
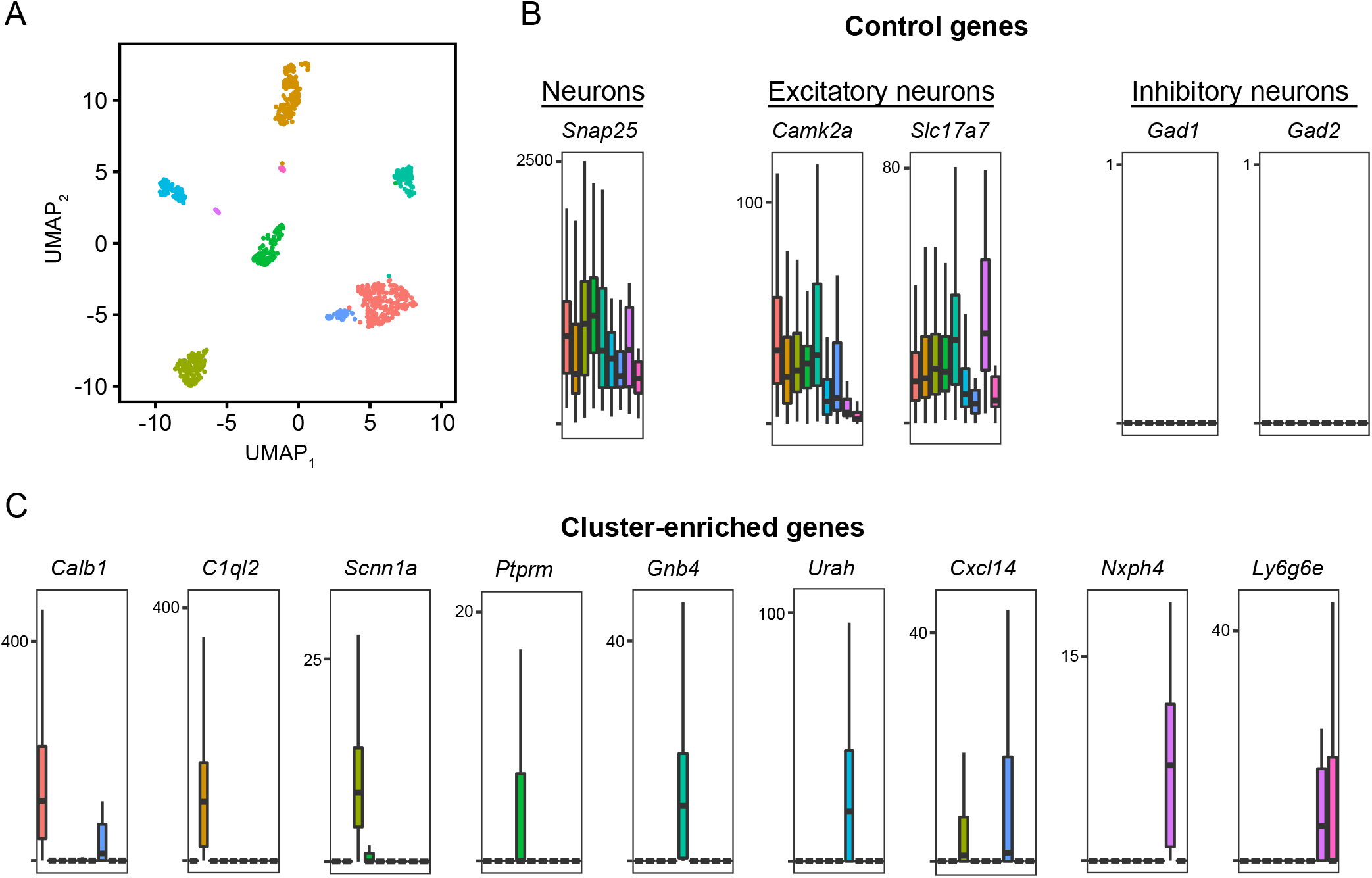
Deconstruction of the RSC excitatory neuron landscape via single-cell RNA sequencing. A. Visualization of single-cell transcriptomes via UMAP nonlinear dimensionality reduction. Cells are coloured according to their respective cluster. B. Boxplots depicting expression of genes for known broad neuron types. C. Expression of cluster-specific markers from scRNA-seq.

Graph-based clustering of this scRNA-seq dataset identified 9 subpopulations, all of which expressed markers of excitatory neurons and lacked expression for markers of interneurons (Figure 1B). These clusters were reproducible across the RSC anterior-posterior axis and across animals (Figure S1B-E), and captured all of the classical laminae of the cortex (Figure S2A). Importantly, individual genes could typically be identified that exhibited expression enriched to single subpopulations (Figure 1C), in principle allowing these subpopulations to be marked and studied by individual genes.

### Validation and spatial registration of RSC cell types

We next sought to map our scRNA-seq subpopulations to spatial locations within the RSC. To first provide a laminar context for our subpopulations, using known neocortical laminar marker genes^13-15^, we analyzed the spatial patterning of RSC laminae via chromogenic *in situ* hybridization from the Allen Mouse Brain Atlas^13^ (Figure S2A,B). Notably, the spatial expression pattens of almost all neocortical laminar markes varied substantially between dRSC and gRSC. *Plxnd4*, a marker of layer 4 granular cells, was expressed only in the gRSC as expected. Expression of *Rgs8*, a marker of layer 2/3 cells, was expressed within dRSC and largely absent in the gRSC. The layer 5 marker *Etv1* showed widespread expression across both the dRSC and gRSC, whereas the layer 6a marker *Foxp2* and the layer 6b marker *Ctgf* primarily showed expression within the dRSC. Of further note, laminar widths and marker gene expression often varied substantially from the neighboring neocortex (Figure S2C), suggesting that the cell-type composition of the RSC might also deviate from neocortex.

Following this examination of cortical laminae in the RSC, we next examined the spatial expression patterns of subpopulation-specific marker genes from our scRNA-seq dataset (Figure 2A). Interestingly, many of these marker genes were enriched within specific laminae, and moreover typically were spatially restricted within either the dRSC or gRSC subregions. For example, the marker genes *Cxcl14, Scnn1a*, and *C1ql2* appeared to be spatially restricted within the gRSC, whereas *Calb1, Rprm*, and *Nxph4* were conversely restricted within the dRSC. Collectively, these genes covered the laminae of the dRSC and gRSC known to contain excitatory neurons (Figure 2B,C), and suggested that there may be discrete changes in cell-type identity within RSC laminae as well as across the dRSC-gRSC border.

**Figure 2.**
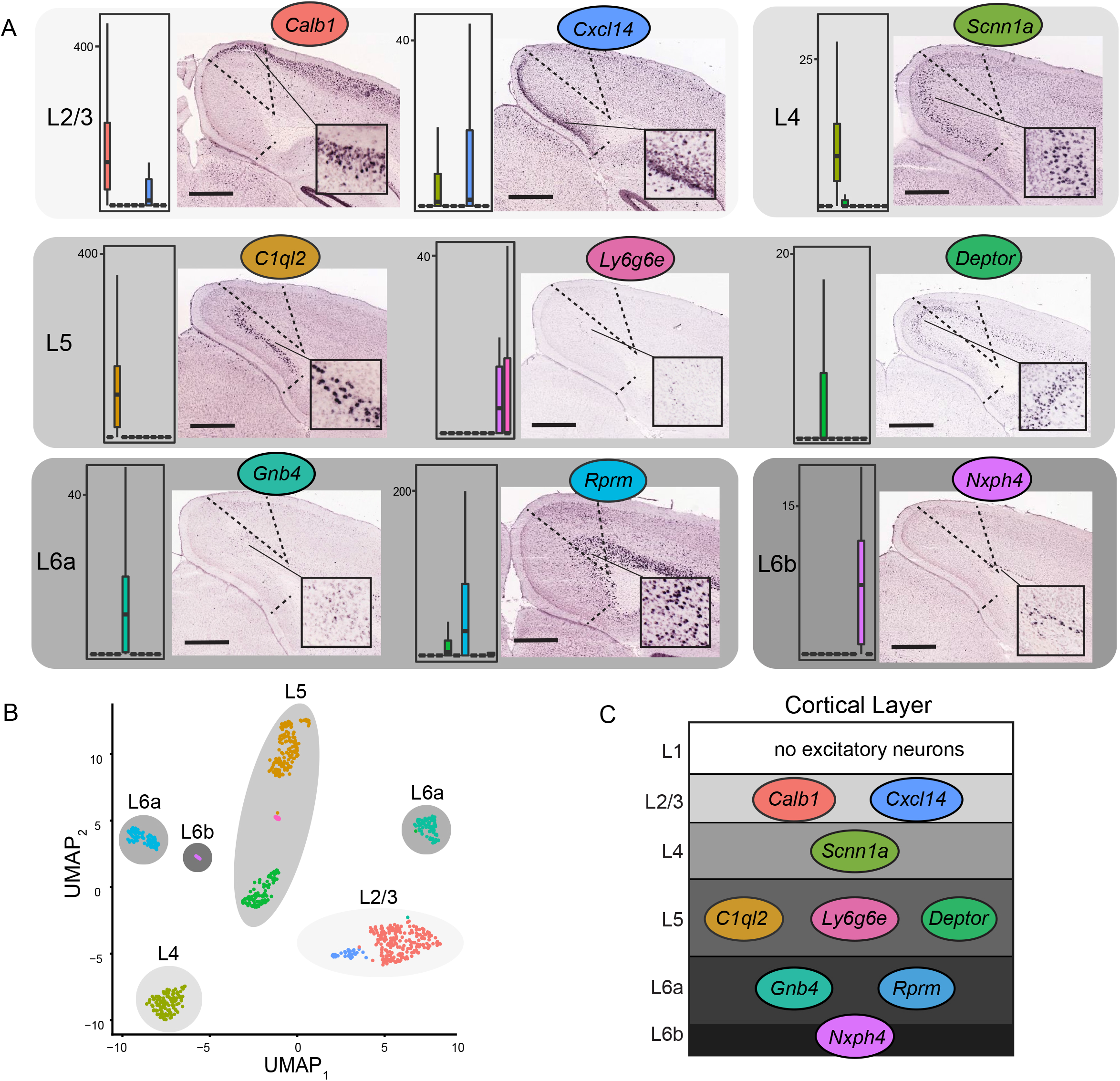
Validation and spatial registration of scRNA-seq marker genes. A. scRNA-seq cluster-specific marker gene expression, as resolved from chromogenic *in situ* hybridization. Scale bar: 839 μm. B. UMAP of scRNA-seq clusters, annotated with putative layer location based upon labeling from (A). C. Schematic illustrating putative spatial location of scRNA-seq clusters.

### Sharp RSC cell-type domains via multiplexed fluorescent in situ hybridization

As the previous chromogenic *in situ* hybridization does not allow direct comparison between multiple genes, we next sought to map all subpopulation marker genes within the same tissue section. We selected marker genes for our scRNA-seq excitatory subpopulations (Figure 1C), and also included genes that identified excitatory neurons broadly (*Slc17a7*) and specific layers for spatial context (layer 2/3: *Calb1*, layer 5: *Etv1*; see Figure S2B). Using these marker genes, we applied a multiplexed single-molecule *in situ* hybridization assay (mFISH), wherein mFISH images were acquired, and *Slc17a7*-expressing excitatory neurons were segmented and analyzed (3407 total *Slc17a7-*expressing putative excitatory neurons were analyzed from 7 sections from 3 animals; see Methods; Figure 3A, S3).

**Figure 3.**
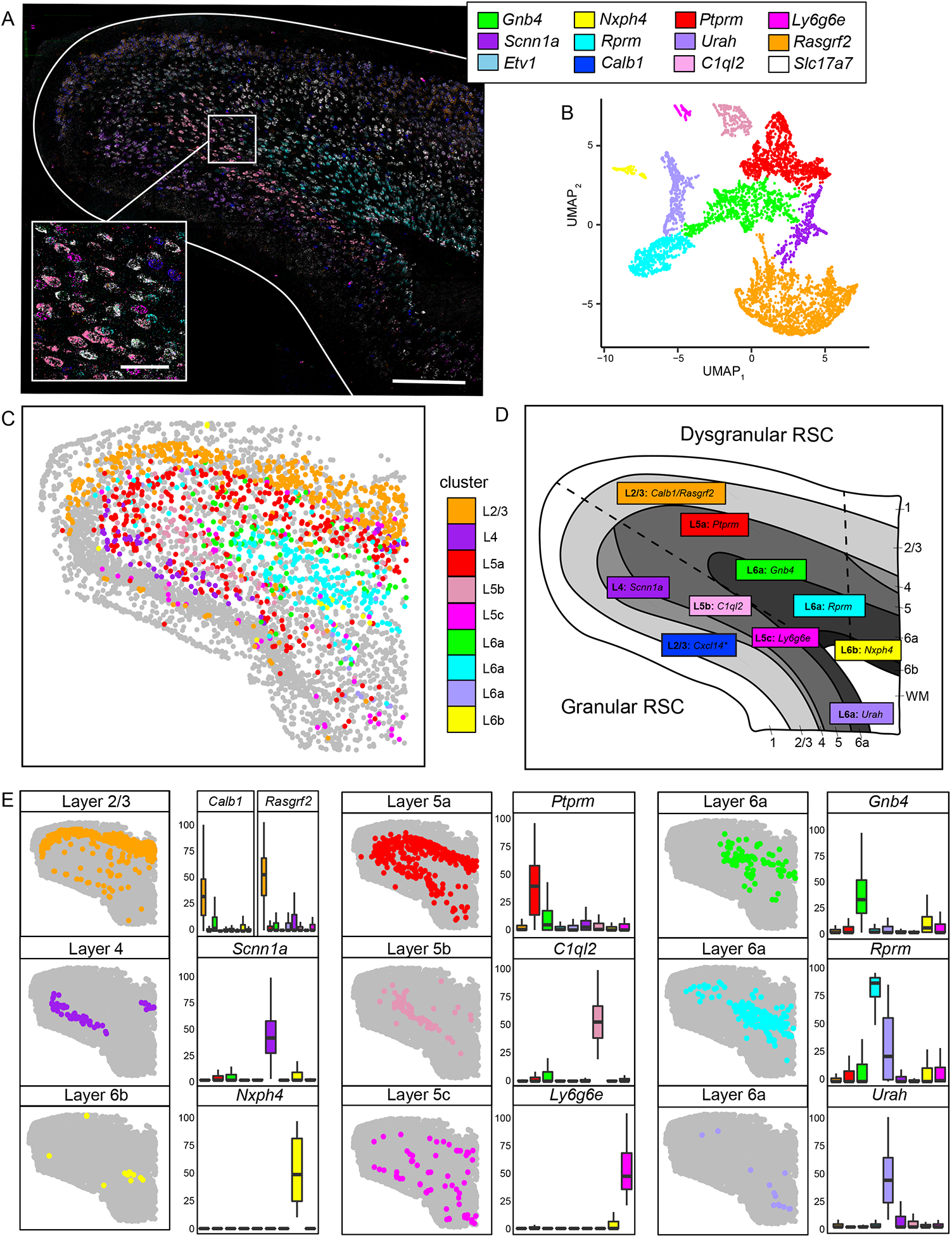
A spatial map of gene expression and cell types using mFISH. A. mFISH image showing posterior RSC with 12 mRNA targets visualized and opaquely overlaid (scale bar: 300 μm). Inset provides expansion (scale bar: 50 μm). Legend denotes the probe and color for the respective target mRNA. B. UMAP embedding of mFISH gene expression data. Cells are coloured according to their cluster identity, with color corresponding to marker gene expression shown in (A). C. Visualization of mFISH clusters in space. Grey points denote non-*Slc17a7* expressing cells. D. Representative schematic displaying cluster identity and marker gene spatial location based on mFISH and chromogenic ISH images. E. Spatial location of all individual excitatory clusters. Boxplots provide expression of marker gene(s) for each cluster.

Clustering (Figure 3B) and spatially mapping (Figure 3C) of *Slc17a7-*expressing neurons revealed the spatial location of excitatory neuronal subpopulations uncovered through scRNA-seq (Figure 3D, E). In particular, each mFISH cluster exhibited localization to a given layer, with many clusters further restricted to the dRSC (*Rprm, Nxph4, Rasgrf2*) or gRSC (*Scnn1a, C1ql2*). These findings recapitulate the marker genes of the scRNA-seq-defined RSC subpopulations, and further demonstrate sharp gene-expression changes between laminae, as well as between the dRSC and gRSC.

### Subregions of the RSC are composed of distinct, spatially restricted cell types

One of the striking findings of our mFISH cell-type mapping was that laminar cell identity was not consistent across the RSC; for example, the marker gene *C1ql2* labelling gRSC layer 5b cells was absent in cells of the dRSC and adjacent neocortex. Motivated by this, we next examined whether gRSC and dRSC subregions could be parcellated by their cell-type composition without requiring *a priori* atlas divisions. To do this, we used a sliding circular window to bin and summarize the quantity of cell types in space, and then hierarchically clustered these binned phenotypes to infer the spatial cell-type-specific organization across the RSC (Figure 4A).

**Figure 4.**
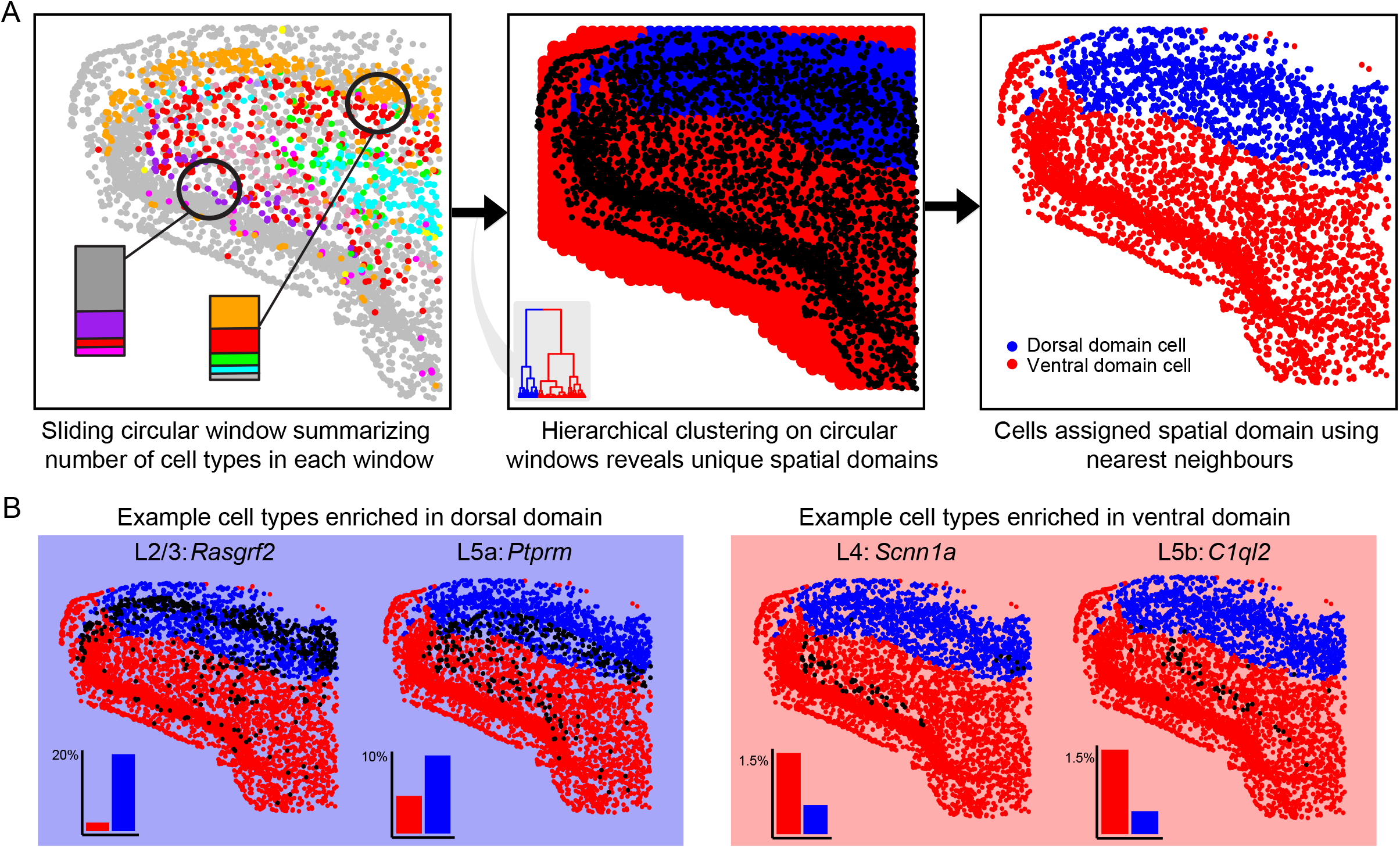
Data-driven RSC subregions derived from cell-type identities. A. A sliding circular window was used to quantify cell types across the tissue section (left), which was clustered to reveal within-RSC spatial domains (middle) and cell types (iii). The blue dorsal cluster captures the upper layers of the dRSC, whereas the red ventral cluster corresponds to the gRSC and layer 6 of the dRSC. B. Example enriched cell-type compositions within the two spatial domains in (A). Insets provide overall percentage of cells within each domain.

Remarkably, the first bifurcation of the dendrogram separated the upper layers of the dRSC from the remaining RSC, illustrating a unique cell type composition for superficial layers of the dRSC. From this bifurcation, we were able to visualize cell-type identities enriched within specific layers of the dRSC (*Calb1, Ptprm*) and the gRSC (*Scnn1a, C1ql2*) (Figure 4B). This finding was reproducible across the anterior-posterior axis of the RSC (Figure S4A,B). This analysis revealed a primary source of within-RSC variation emerging between the dRSC and gRSC, including differences between the putatively homologous layer 2/3 shared between these subregions.

### Cell types vary between the RSC and adjacent visual cortex

Given the large degree of gene-expression changes observed within RSC, we next wanted to examine whether differences also existed between the RSC and the neighbouring visual cortex (VIS). To do this, we downloaded published RSC and VIS single-cell RNA-seq data^9^ and integrated these data with our previous RSC data (see Methods). With this integrated analysis, we performed UMAP embedding and clustering (Figure 5A) and examined whether individual clusters could be identified that were selectively enriched for either RSC or VIS neurons (Figure 5B).

**Figure 5.**
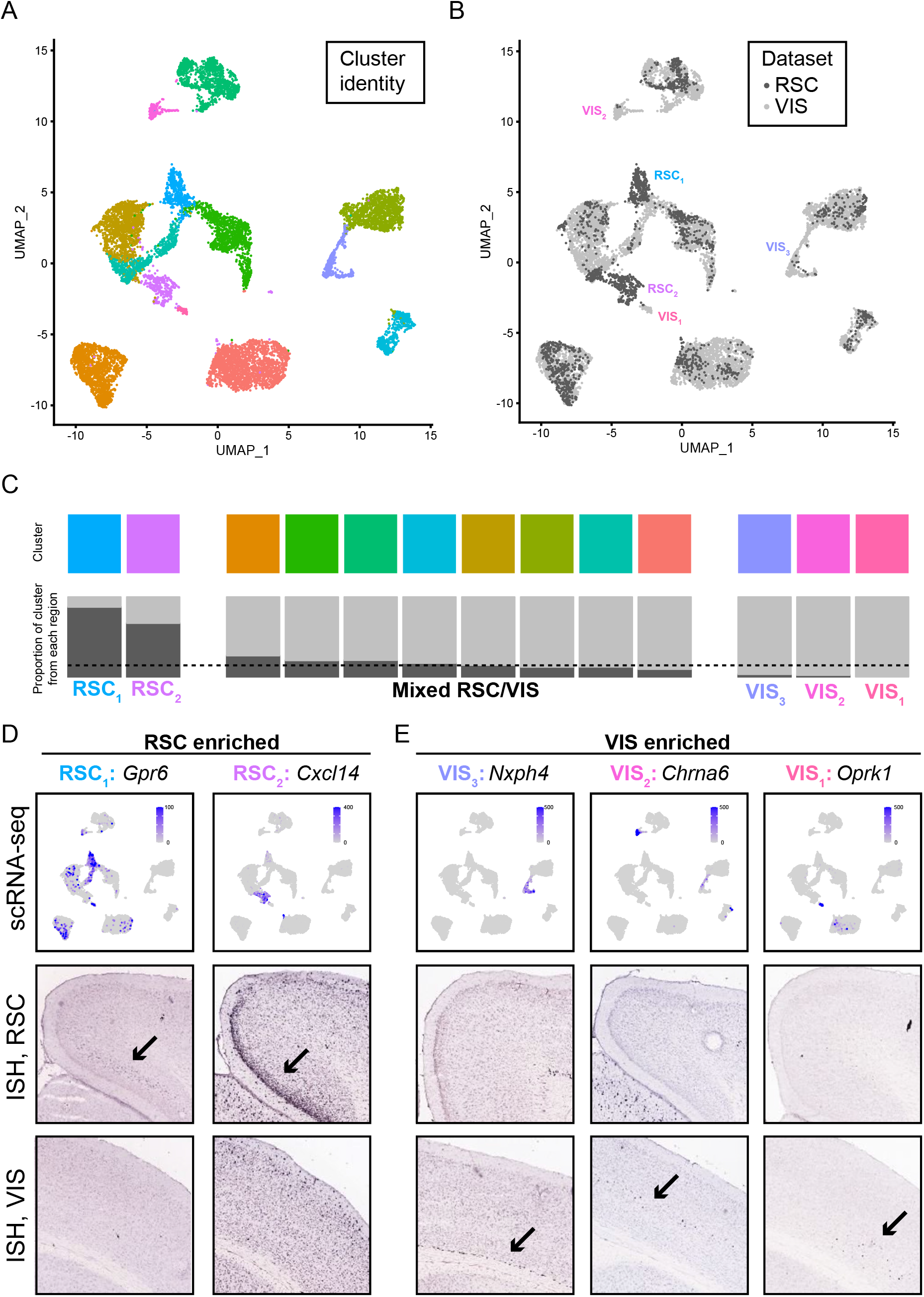
scRNA-seq analysis of RSC vs. VIS reveals region-specific cell types. A. UMAP dimensionality reduction of RSC and VIS datasets, with cells coloured according to cluster. B. As in (A), but cells coloured according to sampled region. C. For each cluster (top row), proportion of cells from each region are shown (bottom row). Dashed line denotes chance levels. D. Expression of individual marker genes for RSC-specific clusters, seen in scRNA-seq UMAP dimensionality reduction (top row) and chromogenic ISH for RSC and VIS (middle, bottom rows). Arrows highlight region of expression. E. As in (D), but for VIS-specific clusters and marker genes.

From this analysis, we were able to identify multiple clusters that were each associated with either RSC or VIS enrichment (Figure 5C). In particular, two clusters enriched selectively for RSC were associated with the marker genes *Gpr6* and *Cxcl14*, which corresponded to layers 4 and 2/3 of the gRSC respectively (Figure 5D; note also that genes for other laminae were RSC-specific: Figure S4C,D). Conversely, the layer 6b marker gene *Nxph4* was enriched in the VIS dataset (Figure 5E, left), as were sparse markers for other neocortical laminae (Figure 5E, middle and right). In collection, these results illustrate excitatory multiple cell types that are not shared between the RSC and the adjacent VIS cortex.

### Sharp cell-type differentiation between the RSC and neocortex

Motivated by our finding of cell-type variation between RSC and VIS, we next investigated whether this RSC-VIS difference generalized to a broader RSC-neocortical difference. To do this, we used a spatial transcriptomics (ST) approach (Visium, 10x Genomics), which uses barcoded spots on tissue slides to spatially map transcripts within tissue sections. We then performed similar dimensionality reduction and clustering analysis to identify the spatial location of transcriptomically similar cortical areas.

Remarkably, we found gRSC-specific spatial clusters for both layer 2/3 and 4, while clusters representing layer 5 and 6 were continuous between RSC and the neighbouring neocortex (Figure 6A,B). These findings were reproducible across animals (Figure S5), and showed specific expression of previous marker genes consistent with our scRNA-seq and mFISH analysis (*Slc17a6* for layer 2/3 or *Scnn1a* for layer 4 of gRSC; Figure 6C). Strikingly, neighboring neocortical regions did not show region-specific clusters for layers 2/3 and 4, but rather exhibited clusters that spanned the length of the neocortical axis. In total, this analysis revealed transcriptional specializations of cell types within the RSC, and the absence of such specializations within other subregions of the neocortex.

**Figure 6.**
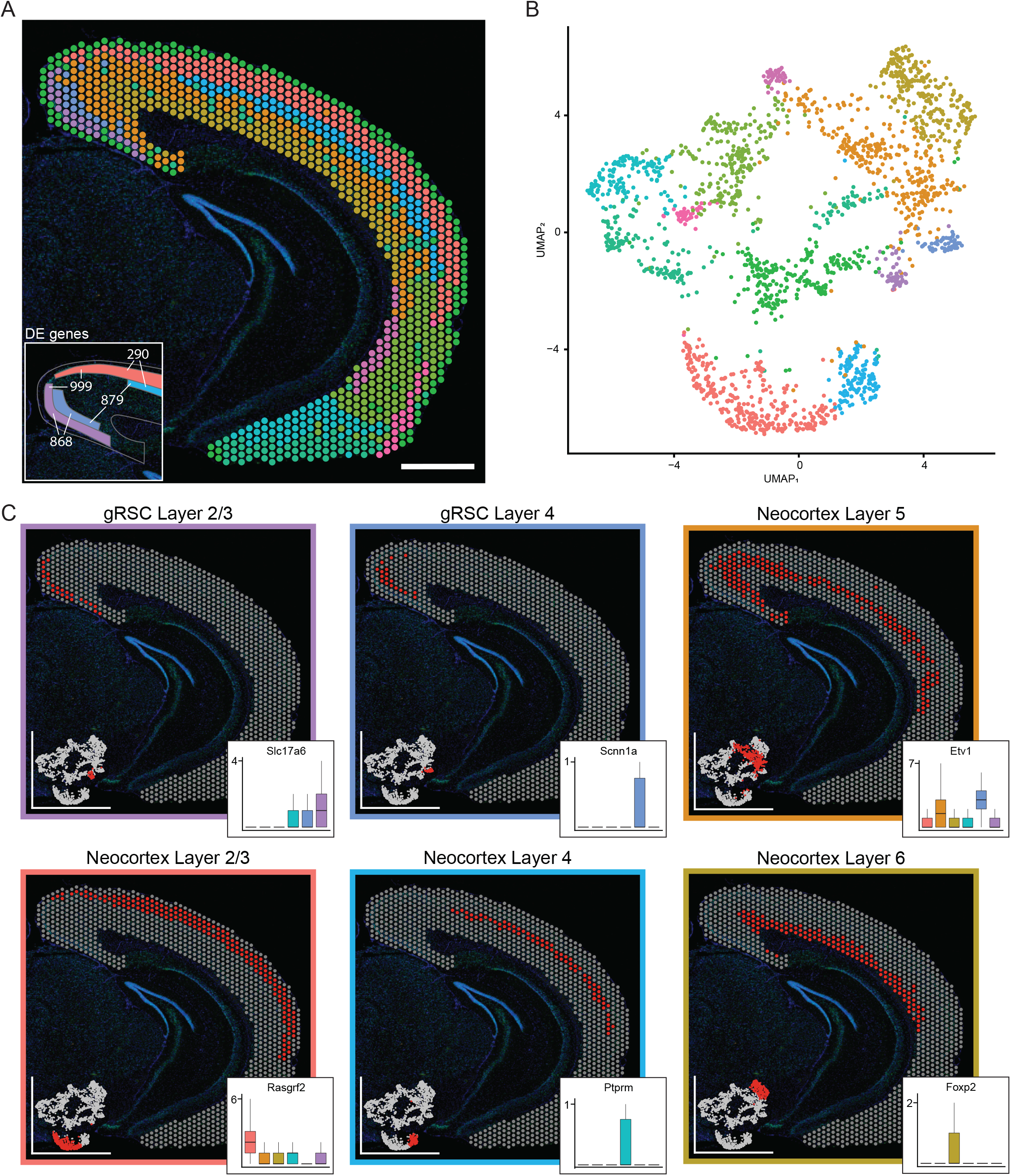
Spatial transcriptomics reveals excitatory cell types unique to the RSC. A,B. Overview of spatial transcriptomics data colored according to cluster identity, as visualized spatially in cortical regions (A) and through UMAP dimensionality reduction (B). Inset in (A) illustrates the number of differentially expressed (DE) genes between pairwise comparisons of clusters of layer 2/3 and 4 within the RSC and neocortex. Scale bar: 1 mm. C. Locations of individual RSC and neocortical clusters from (A), with insets denoting marker genes identified by scRNA-seq and/or mFISH.

The sharp cell-type specializations between layer 2/3 and 4 in the RSC and the neocortex prompted us to quantitatively investigate the extent of transcriptomic differences between these layers and regions. To do this, we examined the total amount of pairwise differentially expressed (DE) genes between these areas (Figure 6A, inset). Remarkably, the highest number of DE genes was present between supposedly homologous laminae of the RSC and neocortex (999 DE genes layer 2/3 gRSC vs neocortex; 879 DE genes layer 4 gRSC vs neocortex). A lesser, but still large, number of DE genes was found between RSC laminae (868 DE genes RSC layer 2/3 vs 4). In contrast, the number of DE genes between neocortical layers was comparatively low (290 DE genes neocortex layer 2/3 vs 4). These relatively large differences between RSC and neocortex laminae highlight the pronounced specialization of RSC cell types, in particular illustrating that their differences relative to neocortical laminar counterparts exceeds the differences associated with distinct laminae.

As ST clustering can potentially occlude gradual spatial changes in cell-type identity, we next performed a ST-cluster-free analysis by correlating our scRNA-seq data (Figures 1, 5) with our ST data (Figure 6). To do this, we extract laminar transcriptomes from either RSC or VIS scRNA-seq (see Methods), and then correlated the expression of the top upregulated genes for each scRNA-seq cluster with individual (i.e. unclustered) ST data points. When correlating with the layer 2/3 gRSC scRNA-seq transcriptome, our results showed spatially restricted high correlation within the gRSC, which dropped sharply into the dRSC and remained low across the neocortex (Figure 7A,B). In contrast, scRNA-seq transcriptomes representing VIS layer 2/3 showed high correlation across millimetres of neocortical regions outside VIS, yet relatively minimal correlation inside gRSC (Figure 7A,B). Such length scales were also seen in layer 4 cell types (Figure 7C,D), and were reproducible across animals (Figure S6). In summary, our data show spatial cluster-independent cell-type differences between RSC and neocortex, and reinforce unique cellular specializations within gRSC layer 2/3 and 4.

**Figure 7.**
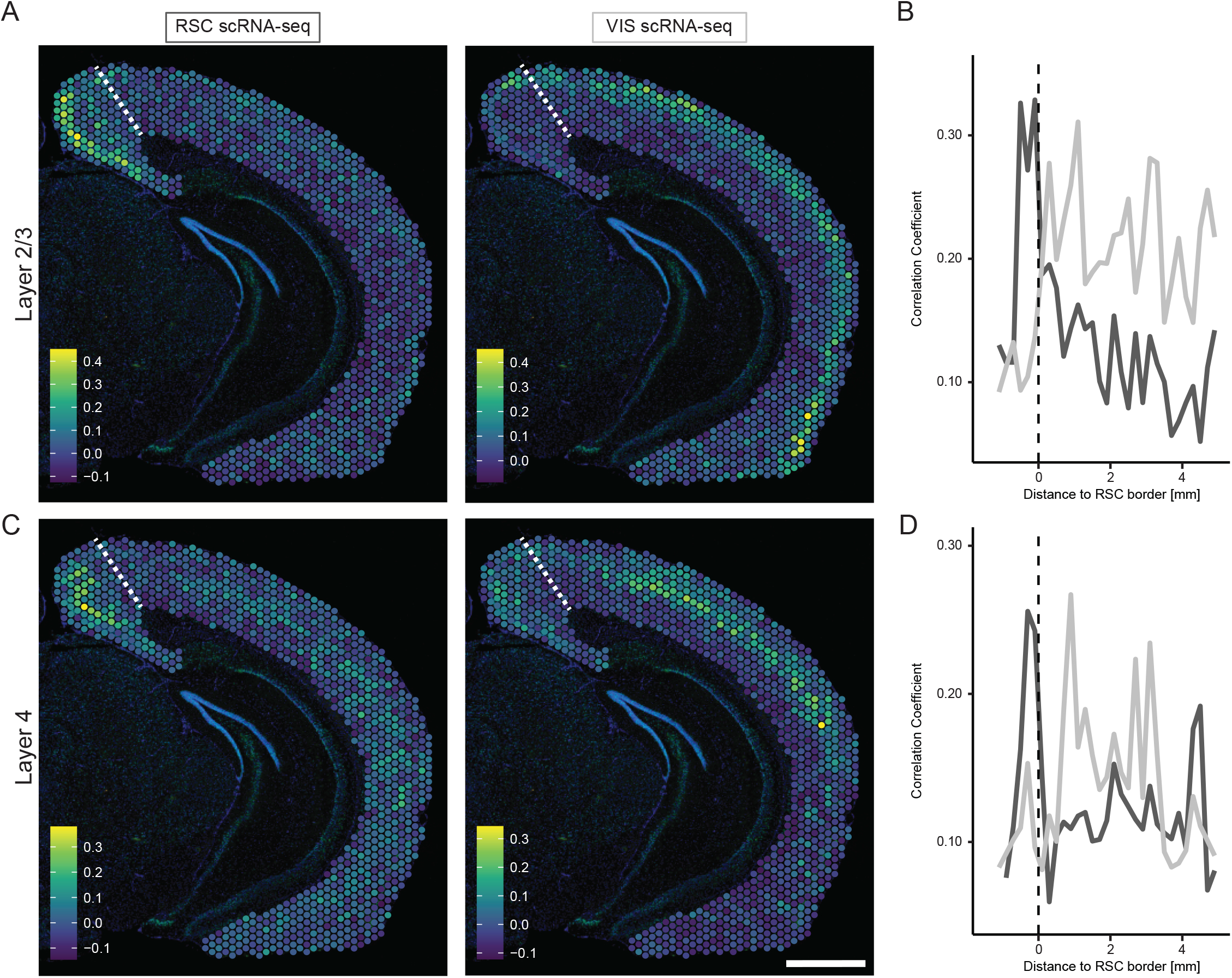
Differential spatial scales of RSC and VIS cell-type identity. A. Pearson correlation coefficient between ST data and average gene expression of scRNA-seq cluster markers for layer 2/3 gRSC (left) or VIS (right). B. Maximum correlation coefficient relative to arc-length distance from RSC border. Dark and light gray denote correlation of ST data with gRSC and VIS, respectively. C,D. As in (A,B), but for layer 4 marker genes. Scale bar: 1 mm.

## DISCUSSION

Neocortical laminae are posited to be a relatively stereotyped across brain regions, with changes to cell-type organization and other properties typically occurring in a graded fashion across multiple regions. To examine this assumption, here we investigated whether cell-type differences were present within distinct regions of the RSC, as well as in the RSC relative to the neocortex. In combining scRNA-seq, mFISH, and spatial transcriptomics, we identified multiple spatially restricted and transcriptomically unique excitatory neuron cell types within the RSC. These findings have important implications in interpreting RSC structure and function, and provide future guidance into the organizational rules and functional mechanisms of different cortical regions.

### Phenomenology and mechanisms of differential cortical organization

In previous literature, two seemingly disparate views of the cortex have been established^16^. From the perspective of cell-type identity, the cortex is often considered a relatively homogenous structure^2^, with changes in gene expression and constituent cell types occurring gradually over long distances^17^. From a functional perspective, the cortex is conversely typically viewed as a patchwork of functionally discrete areas with spatially sharp borders^18^, such as the cytoarchitectonically defined Brodmann areas^19^. Here, our findings illustrate that distinct brain regions can simultaneously capture both of these perspectives. In particular, whereas the RSC exhibits specialized cell types that are restricted and discrete in space, cell-type identity in the neighbouring neocortex remains relatively homogenous along its length (Figures 6, 7).

These differences in organizational principles between the RSC and the neocortex may be associated with the phylogenetic age and spatial location of these regions. Traditionally, neuroanatomists have divided the cortical plate into the 6-layered isocortex (neocortex) and the 3-4 layered allocortex (comprising areas such as the subiculum, the hippocampal formation, and the entorhinal cortex)^20^. These allocortical regions are noted to have more cross-region heterogeneity than their isocortical counterparts, and also thought to be phylogenetically older in age^21,22^. As the RSC is spatially adjacent to the allocortical regions of the postsubiculum and dorsal subiculum, it may be that the RSC is effectively a transitionary region between the rodent allocortex and neocortex. Consistent with this, our ST findings show that other allocortical regions also exhibit sharp changes in transcriptomic identity and layer composition (e.g., the entorhinal cortex: Figure 6). Thus, one explanation for the unique layer composition of the RSC is an older phylogenetic age, which manifests as unique cell-type compositions and organizational principles relative to the neocortex.

### Complementary molecular methodologies converge on unique RSC cell types

Here, our findings demonstrate specialized properties of RSC cell types, wherein gRSC layer 2/3 and layer 4 neurons are markedly different relative to neocortical counterparts (Figures 5,6,7). Such a cell-type specialization appears particular to the RSC, rather than a general feature of regional specializations across the cortex, as our ST data did not further differentiate between neocortical regions (e.g., between the visual cortex, posterior parietal cortex, and auditory cortex: Figure 6). Although ST-cluster-based differentiation between VIS and other neocortical areas could in principle miss sparse cell populations, VIS-enriched genes were generally expressed across multiple cortical areas (Figure 5E), with correlations spanning millimeters of neocortex (Figure 7). As such, it appears that neocortical laminar cell-type identities are relatively stereotyped, which is in agreement with current findings and can be directly verified by other existing datasets^23^ in future work. Ultimately, our results highlight that the distinct cell-types of the RSC are unique in their heterogeneity relative to entire neocortex.

### Importance of understanding the cell types of the RSC

The retrosplenial cortex is frequently studied *in vivo* due to its experimentally convenient location at dorsal midline^24-27^. As a consequence, understanding the cell-type identities underlying these *in vivo* recordings is of particular importance. Here, we used a data-driven approach to parse out layers (Figure 3) and subregions (Figure 4) of the RSC at a single-cell resolution, providing geometrical constraints on the location and identities of excitatory RSC cell types. Our results demonstrate a gRSC/dRSC specificity in cell type composition, as well as uncover differences between the gRSC and the neocortex (Figure 6). This may provide insight into clarifying functional differences between the gRSC and dRSC^10,28-31^, as well as revealing differences in RSC information processing relative to the neocortex.

Our work also informs future experimental avenues and interpretation. Methodologically, the transcriptomic profiles of these RSC cell types can be leveraged to identify relevant genes for attaining genetic access to a specific cell type (e.g., via Cre-mediated transgenic mice lines), as well as to investigate causal roles of individual genes (eg: via CRISPR-Cas9 gene deletion^32^). Our work also emphasizes that the cell-type differences between supposedly homologous laminae may provide interpretational constraints to experiments. In particular, *in vivo* imaging techniques (e.g. calcium imaging^33^) and interventional techniques (e.g. DREADDs^34^) rely on cells with stereotyped intrinsic mechanisms and signaling cascades, which are violated when examining neurons with variable cell-type identity. Ultimately, our data provides access to each cell types transcriptome, allowing one to refine experimental procedures and reproducibly target cells and molecules, while avoiding methodological caveats.

### Differential computations within RSC subregions

Our results demonstrate changes in cell type composition RSC laminae and subregions, which may underlie known distinct functional properties of individual cells in the RSC. For example, two electrophysiologically unique excitatory cell types have been reported in layer 2/3 of the gRSC. One is a low rheobase cell type with unique dendritic morphology (matching spatially with the *Cxcl14*-expressing cell type identified here), whereas the other is a regular spiking cell type (which matches spatially to the *Rasgrf2*-expressing cell type identified here)^35^. This illustrates that some of our identified cell types may have unique electrophysiological properties, which can be targeted and examined in future work. Although comprehensive investigation of single-cell properties across the dRSC/gRSC border have yet to be performed, the large transcriptomic differences identified here likely suggest similar functional differences along this spatial axis.

The cell type composition of the gRSC and dRSC may further underlie their reported functional differences at a behavioural level. Broadly, the RSC is known to contain head-direction cells^10^, complex conjunctive coding cells^11^, and fear memory engram cells^6^, which are differentially distributed within RSC subregions. In particular, the dRSC is implicated more in head direction, place, and velocity tuning^10,11^, whereas the gRSC has been reported to contain fear memory engrams for both recent and remote memory^6^. At the level of cell-type specificity, ablation of layers 4 and 5 of the gRSC (corresponding to the *Scnn1a*-expressing and *C1ql2*-expressing cell types identified here) causes both retrograde and anterograde amnesia in rats^36^. This suggests the differences in dRSC and gRSC cell type composition might contribute to these differences in layer and subregion-specific behaviours.

Ultimately, the cell-type differences identified here likely result in very different local network computations within dRSC and gRSC relative to the neocortex. The canonical neocortical circuit receives subcortical afferents to layer 4, which is transmitted to layer 2/3 and combined with corticocortical inputs. These signals are then sent routed to layers 5 and 6, which ultimately send efferents to subcortical and thalamic targets respectively^37^. Notably, dRSC lacks a layer 4 (Figure 3), meaning that a fundamental processing layer for receiving subcortical input is absent in the dRSC relative to the neocortex. Although the gRSC contains a layer 4, both this layer and layer 2/3 are markedly different in intrinsic cell-type identify relative to the neocortex (Figures 6,7). These marked differences, for both the dRSC and gRSC relative to the neocortex, strongly suggest differential processing of subcortical and corticocortical information in the RSC.

### Prospective insight from the cell-type organization of the RSC

Our findings allow examination of the roles of well-defined RSC excitatory neurons with cell type, layer, and subregion specificity (Figures 1,2,3). Importantly, understanding the RSC at these granularities will provide cell-type and spatial frameworks where experimental results can be interpreted and interrelated. As one example at the intersection of all of these granularities, we have identified multiple cell types with unique transcriptomic profiles that are spatially restricted to sublaminae of layer 5 RSC subregions (*Ly6g6e-* and *C1ql2-*expressing neurons: Figure 4D, Figure S4C). Such cells, identifiable by both marker genes and spatial locations, can be recognized, labeled, and manipulated across an array of future experiments to assess their cell-type-specific contributions. More broadly, our findings reveal distinct differences in the layers 2/3 and 4 of the gRSC compared to the neocortex, which along with our online resource (http://scrnaseq.janelia.org/rsc), will allow many potential avenues for future research. As such, this framework of cell types in the RSC provided here will help guide correlative and causal interpretation of specialized genes, cells, circuits of the RSC.

## METHODS

Unless otherwise noted, experiments used male wild-type C57BL/6 mice between 8-20 weeks of age. All experimental procedures were approved by the University of British Columbia Animal Care Committee and the Janelia Research Campus Institutional Animal Care and Use Committee.

### scRNA-seq data generation

To generate scRNA-seq data for analysis, we proceeded according to published methods ^12,38^. Specifically, in n=3 C57bl/6 mice, manual purification ^39^ was used to capture cells in capillary needles in approximately 0.1-0.5 mL ACSF cocktail, placed into 8-well strips containing 3 μL of cell collection buffer (0.1% Triton X-100, 0.2 U/μL RNAse inhibitor (Lucigen)), and generally processed according to published methodology ^40,41^. Specifically, each strip of cells was flash frozen on dry ice, then stored at −80°C until cDNA synthesis. Cells were lysed by adding 1 μL lysis mix (50 mM Tris pH 8.0, 5 mM EDTA pH 8.0, 10 mM DTT, 1% Tween-20, 1% Triton X-100, 0.1 g/L Proteinase K (Roche), 2.5 mM dNTPs (Takara), and ERCC Mix 1 (Thermo Fisher) diluted to 1e-6 and 1 μL 10 μM barcoded RT E3V6NEXT primer^42^. The primer was modified to add a 1 bp spacer before the barcode, extending the barcode length from 6bp to 8bp, and designing the 384 barcodes to tolerate 1 mismatch error correction. The samples were incubated for 5 minutes at 50°C to lyse the cells, followed by 20 minutes at 80°C to inactivate the Proteinase K. Reverse transcription master mix (2 μL 5X buffer (Thermo Fisher Scientific), 2 μL 5M Betaine (Sigma-Aldrich), 0.2 μL 50 μM E5V6NEXT template switch oligo (Integrated DNA Technologies) ^42^, 0.1 μL 200 U/μL Maxima H-RT (Thermo Fisher Scientific), 0.1 μL 40 U/μL RNasin (Lucigen), and 0.6 μL nuclease-free water (Thermo Fisher Scientific) was added to the approximately 5.5 μL lysis reaction and incubated at 42°C for 1.5 hour, followed by 10 minutes at 75°C to inactivate reverse transcriptase. PCR was performed by adding 10 μL 2X HiFi PCR mix (Kapa Biosystems) and 0.5 μl 60 μM SINGV6 primer with the following conditions: 98°C for 3 minutes, 20 cycles of 98°C for 20 seconds, 64°C for 15 seconds, 72°C for 4 minutes, with a final extension step of 5 minutes at 72°C. Samples were pooled across the plate to yield approximately 2 mL pooled PCR reaction. From this, 500 μL was purified with 300 μL Ampure XP beads (0.6x ratio; Beckman CoulterA), washed twice with 75% ethanol, and eluted in 20 μL nuclease-free water. The cDNA concentration was determined using Qubit High-Sensitivity DNA kit (Thermo Fisher Scientific). Eleven plates were analyzed in total, with six hundred pg cDNA from each plate of cells used in a modified Nextera XT (Illumina) library preparation with 5 μM P5NEXTPT5 primer ^42^. The resulting libraries were purified according to the Nextera XT protocol (0.6x ratio) and quantified by qPCR using Kapa Library Quantification (Kapa Biosystems). Three plates were pooled on a NextSeq 550 mid-output flowcell with 26 bases in read 1, 8 bases for the i7 index, and 125 bases in read 2, and the remaining eight plates were pooled on a NextSeq 550 high-output flowcell with 26 bases in read 1, 8 bases for the i7 index, and 50 bases for read 2. Alignment and count-based quantification of single-cell data was performed by removing adapters, tagging transcript reads to barcodes and UMIs, and aligned to the mouse genome via STAR ^43^.

In total, 907 cells were manually harvested and underwent sequencing. Of these initial 907 cells, 765 we retained for analysis after filtering against non-neuronal cells (*Snap25* CPM>100), oligodendrocytes (*Olig2* CPM =0), and interneurons *(Gad1* CPM < 10) and creating a Seurat object via *CreateSeuratObject()* with *min*.*cells=10* and *min*.*features=2500*. These 765 cells exhibited 5.4 ± 1.0 thousand expressed genes/cell from 138 ± 84 thousand reads/cell (mean ± SD). The relatively high abundance of excitatory neurons sampled owed to the fact that excitatory neurons are relatively abundant relative to interneurons in the retrosplenial cortex. No blinding or randomization was used for the construction or analysis of this dataset. No *a priori* sample size was determined for the number of animals or cells to use; note that previous methods have indicated that several hundred cells from a single animal is sufficient to resolve heterogeneity within excitatory neuronal cell types ^41,44^.

Computational analysis was performed in R (RRID:SCR_001905) ^45^ using a combination of Seurat v4 (RRID:SCR_007322) ^46,47^ and custom scripts ^41^. To analyze our data, the Seurat object was processed via *SCTransform(), ScaleData(), RunPCA(), RunTSNE(), FindNeighbors(k = 12,dims=1:15), FindClusters(resolution=0*.*7), RunUMAP(reduction=‘pca’)*, with all parameters used as default values unless otherwise listed. This processed Seurat object was then used for downstream analysis. Cluster-specific enriched genes obeying *p*_*ADJ*_ *<* 0.05 were obtained with Seurat via *FindMarkers()*, where *p*_*ADJ*_ is the adjusted *p*-value from Seurat based on Bonferroni correction. For boxplot visualizations, values from the Seurat data slot are shown, with hinges denoting first and third quartile and whiskers denoting remaining data points up to at most 1.5 * interquartile range. Outlier values beyond whiskers are not shown. Raw and processed scRNA-seq datasets have been deposited in the National Center for Biotechnology Information (NCBI) Gene Expression Omnibus under GEO: GSE200027.

### Integration with published single-cell RNA sequencing data

To integrate and compare our scRNA-seq data to previously published data, we downloaded RSC and visual cortex data from a recent publication^23^, using the identifiers “RSP” and “VIS” from this dataset. Thresholding and Seurat object creation were performed analogously to original RSC data. Each individual dataset underwent an SCTransform operation (via *scSeurat*.*list <-SplitObject(split*.*by=‘dataset’); scSeurat*.*list <-lapply(scSeurat*.*list, SCTransform); scSeurat*.*features <-SelectIntegrationFeatures(scSeurat*.*list, nfeatures=3000, verbose=F); scSeurat*.*list <-PrepSCTIntegration(object*.*list=scSeurat*.*list, anchor*.*features=scSeurat*.*features); int*.*anchors <-FindIntegrationAnchors(object*.*list=scSeurat*.*list, normalization*.*method=‘SCT’, anchor. features=scSeurat*.*features); scSeurat*.*integrated <-IntegrateData(anchorset=int*.*anchors, normalization*.*method=‘SCT’, verbose=F)*. Subsequent analysis on the integrated dataset, starting with *ScaleData()*, was handled analogously to the original RSC dataset, with *dims=1:20, resolution = 0*.*25*, and all other parameters being default. This analysis involved a total of 10,978 cells, with 10.0 ± 1.9 thousand expressed genes/cell from 1546 ± 768 thousand reads/cell (mean ± SD), with 80.0% of the dataset being from VIS (n=8787 cells) and 20.0% from pooled RSC (n=2191 cells). Further computational analysis, included obtaining marker genes, was handled analogously to the original RSC dataset.

### mFISH data acquisition and analysis

The following custom probes from Advanced Cell Diagnostics were purchased for mFISH: *Gnb4* (Cat No. 460951-T1), *Ly6g6e* (Cat No. 506391-T2), *Ptprm* (Cat No. 579201-T3), *Nxph4* (Cat No. 489641-T4), *Scnn1a* (441391-T5), *Rprm* (Cat No. 466071-T6), *Urah* (Cat No. 525331-T7), *Rasgrf2* (Cat No. 467881-T8), *Etv1* (Cat No. 557891-T9), *Calb1* (Cat No. 428431-T10), *C1ql2* (Cat No. 480871-T11), and *Slc17a7* (Cat No. 501101-T12). mFISH was implemented procedurally in the same manner as previously published^12,38^. Briefly, mature male mice were randomly selected for mFISH and were deeply anesthetized with isoflurane and perfused with phosphate buffered saline (PBS) followed by 4% paraformaldehyde (PFA) in PBS. Brains were dissected and post-fixed in 4% PFA for 24 hrs, then cryoprotected in a solution of 30% sucrose in PBS for 48 hrs. Brain sections (20 μm) were made using a cryostat tissue slicer and mounted on coated glass slides. Slides were subsequently stored at –80°C until use.

For use, the tissue underwent pre-treatment and antigen retrieval per the User Manual for Fixed Frozen Tissue (Advanced Cell Diagnostics). All 12 probes with unique tails (T1–T12) were hybridized to the tissue, amplified, and the tissue counterstained with DAPI. Probes were visualized four at a time using cleavable fluorophores in the 488nm, 550nm, 647nm, and 750nm channels. This occurred via an iterative process of imaging, de-coverslipping, fluorophore cleaving, and adding the next four targeted fluorophores, until all 12 gene-expression targets were imaged.

mFISH images were acquired with a 63× objective on a SP8 Leica white light laser confocal microscope (Leica Microsystems). Z-stacks were acquired with a step size of 0.45 μm for each imaging round. Final composite images are pseudocolored maximum intensity projections, with channels opaquely overlaid upon one another ordered from highest to lowest expression. Presented images include brightness adjustments applied to individual channels uniformly across the entire image, and a linear smoothing filter utilized on probes in the 750nm channel (*Nxph4*-T4, *Rasgrf2*-T8, *Slc17a7*-T12) to accommodate noise introduced via increased gain and laser power required in this channel during imaging.

Processing of mFISH images generally followed our previously published analysis pipeline^12,38^ using Fiji^48^. Briefly, maximum intensity projections of the DAPI signal from each round was used to rigidly register probe signals across rounds, followed by nonlinear elastic registration via bUnwarpJ^49^ to accommodate any nonlinear tissue warping due to de-coverslipping. The individual nuclei from each round were then binarized and multiplied to include only cells present across all three rounds. These were then segmented and dilated by 3 μm to include the surrounding cytosol. The signal from each probe was then binarized by thresholding at the last 0.2–1% of the histogram tail, and then the number of pixels within regions of interest (ROIs) selected from segmentation was summed and normalized to the pixel area of the cell and multiplied by 100. This in effect corresponded to percent area covered (PAC) of the optical space of a cell.

Three mature male mice with sections from the anterior, intermediate, and posterior RSC were utilized, with a total of 7 sections undergoing mFISH. Across these three animals, 15,553 cells in total were imaged. To facilitate analysis of excitatory neurons specifically, a threshold of 0.5 PAC of *Slc17a7* was required for each cell to be included in analysis, resulting in 6,662 total putative excitatory neurons being used for subsequent analysis. *Slc17a7* expression levels were used only for cellular phenotyping, and thus excluded from further analysis. PAC matrices were dimensionally reduced via UMAP^50^ using a Manhattan metric with nearest-neighbors value of 15 and a minimum distance of 0.09, and cells were clustered via Leiden clustering^51^ from the Monocle 3^52^ package with a resolution of 4×10^−4^.

For unbiased analysis of cell-type clustering (Figure 4), a sliding circular window of radius 250 μm and a step size of 30 μm were used to aggregate the number of cell types in a given region (akin to previously published work^53^). The data was then subset to exclude windows with a density of cells less than 0.4 and hierarchical clustering was performed. In order to identify differentially enriched cell types in space, the dendrogram was cut at the first bifurcation. Boxplots for the spatial domain analysis were calculated by taking the percentage of a given cell type (*Ptprm, Rasgrf2, C1ql2, Scnn1a)* within either the dorsal or ventral spatial domains, relative to the total number of cells within each domain.

### Spatial transcriptomics data acquisition and analysis

For Visium ST analysis, two mice were deeply anesthetized with isoflurane, decapitated and brains were quickly removed, embedded in OCT and frozen in methylbutane cooled down using liquid nitrogen. Processing of slices and library preparation were performed for Visium as recommended by the manufacturer (10x Genomics). In brief, 10 um slices of posterior RSC were placed on 10x genomics Visium slides, fixed with methanol and labeled using NeuN 488 conjugated antibody (1:100, Abcam, RRID: AB_2716282) and DAPI (0.2 mg/ml, Sigma). Images were acquired using a 10x objective on a Zeiss LSM 880 confocal microscope. Images included fiducial frame in TRITC for spot alignment in downstream analysis. Slices were permeabilized for 24 minutes (previously optimized using samples of same genotype and age) and libraries were prepared according to the manufacturer’s user guide. Libraries were sequenced using an Illumina NextSeq2000 at a reading depth of 100M reads per sample using the following read metrics: Read1: 28bp; Index1: 10bp; Index2: 10bp; Read 2: 90bp. Raw FASTQ files were processed with Space Ranger 1.3.0 and aligned to mm10-2020-A reference genome (refdata-gex-mm10-2020-A). Manual fiducial alignment and selection of spots under cortex was performed in Loupe browser 6. ST data was analyzed using Seurat package (version 4) in R (version 4.1.2) and in-house written code. Data of both slices was normalized (via *SCtransform*), filtered to remove mitochondrial genes and integrated using Seurat’s anchored integration. Using principal component analysis, we identified the first 15 PCs for downstream analysis. Data was then clustered and plotted using UMAP non-linear dimensionality reduction (*resolution=0*.*6*).

For correlation analysis between scRNA-seq data and ST data we computed the average gene expression of the top 50 upregulated genes identified using the function *FindAllMarker(only*.*pos = TRUE, min*.*pct = 0*.*25, logfc*.*threshold = 0*.*25)* for each scRNA-seq cluster. Genes for all clusters (RSC and VIS separately) were combined and used to calculate Pearson correlation between each cluster and each spot of ST data for all slices. We found using a subset of cluster defining genes (max 50 per cluster, totaling 313 genes for RSC and 376 for VIS) enabled us to show discrete differences between cortical regions since the majority of expressed genes are likely shared between similar cell types of different regions.

Correlation coefficient was plotted in space over cortex using spot coordinates assigned during manual image alignment in Loupe browser. Correlations were plotted over arc-length distance from border between RSC and neighboring cortex. Since the RSC border was defined as a line perpendicular to the pial surface, spots with the same distance (calculated using Manhattan distance) to this border have the same orientation. By calculating maximum correlation values for each distance, we were able to plot the correlation for different regions of the cortex along the layers of the cortex.

### Statistical conventions

Unless otherwise noted, the following conventions were used in this study. Box-and-whisker plots show distribution of gene expression within cell populations according to the following conventions: midline denotes median, boxes denote first and third quartiles, whiskers denote remaining data points up to at most 1.5 * interquartile range, with outlier values beyond whiskers not shown. *p*_*ADJ*_*-*values are computed via a Mann-Whitney U test with FDR correction for multiple comparisons. Statistical significance for adjusted *p*_*ADJ*_-values is denoted as: ns: *p*_*ADJ*≥_0.05; *: *p*_*ADJ*_<0.05, **: *p*_*ADJ*_<0.01, ***: *p*_*ADJ*_<0.001.

## ACKNOWLEDGEMENTS

MSC is supported by the University of British Columbia (Department of Cellular and Physiological Sciences, Djavad Mowafaghian Centre for Brain Health, and the Faculty of Medicine Research Office), the Natural Sciences and Engineering Research Council of Canada (RGPIN-2019-04507), the Canadian Institutes of Health Research (PJT-419798), and the Canadian Foundation for Innovation (John R. Evans Leaders Fund 38369). KES is supported by a Royal Canadian Legion Masters Scholarship in Veteran Health Research from the Canadian Institute for Military and Veteran Health Research, a Canada Graduate Scholarship – Masters from the Canadian Institute of Health Research, and a Canada Graduate Scholarship – Doctoral from the Natural Sciences and Engineering Research Council. LK is funded by the Deutsche Forschungsgemeinschaft (DFG, German Research Foundation, Walter Benjamin fellowship, project 444112617). This work was supported by resources made available through the NeuroImaging and NeuroComputation Centre at the Djavad Mowafaghian Centre for Brain Health (RRID: SCR_019086). Collaboration between MSC and LW, JC, and ALL was supported by the Janelia Visiting Scientist Program. We thank members of the Cembrowski lab for helpful discussions, and Jeffrey LeDue for insight and guidance in image acquisition.

## SUPPLEMENTAL FIGURES

**Figure S1. Consistency of single-cell transcriptomes across spatial locations and animals**. Related to Figure 1.

A. Schematized location of microdissected anterior, intermediate, and posterior locations. B. UMAP dimensionality reduction showing microdissected region for each cell. C. As in (B), but showing original animal for each cell. D. UMAP dimensionality reduction showing cluster identity, for comparison to (B,C). E. Quantification of number of cells for each cluster.

**Figure S2. Spatial mapping of layer-specific marker genes in the RSC and neocortex**.

Related to Figure 2.

A. Expression of laminar marker genes in scRNA-seq RSC dataset. B. Expression of laminar marker genes in chromogenic ISH dataset in RSC. Scale bar: 839μm. C. As in (B), but for region V1 to compare layer-specific labeling to RSC. Individual rows illustrate marker genes for different cortical laminae. Scale bar: 419μm.

**Figure S3. mFISH clusters and marker genes across the anterior-posterior axis of the RSC**.

Related to Figure 3.

A. mFISH images of anterior and intermediate RSC (scale bar: 300 μm). Legend provides marker genes and coloring conventions, and insets provide expansion of boxed area (scale bar: 50 μm). B. Clustering of mFISH data for anterior and intermediate sections. Inset provides UMAP for reference. C. Individual clusters in space, with boxplots providing marker gene expression for each cluster.

**Figure S4. Subregions across the anterior-posterior axis of the RSC**.

Related to Figure 4.

A. Spatial domain analysis of the anterior RSC, with hierarchical clustering and cell type composition analyses as in Figure 4. B. As in (A), but for the intermediate RSC. C. Expression of *C1ql2*, an RSC-specific marker gene of L5b, across the anterior-posterior axis of the RSC. Image from the Allen Mouse Brain Atlas. D. As in (C), but for the RSC-restricted L5c marker gene *Ly6g6e*.

**Figure S5. Reproducibility of spatial transcriptomics clusters**.

Related to Figure 6.

A,B. Spatial transcriptomics data colored according to cluster identity for 2 replicates (A and B, respectively). Scale bar: 1 mm. C. Individual clusters highlighted in red spatially shown for two replicates (top and middle rows, respectively), with UMAP dimensionality reduction provided for reference (bottom row).

**Figure S6. Reproducibility of region-specific correlation of ST and scRNA-seq**.

Related to Figure 7.

A. Pearson correlation coefficient between ST data and average gene expression of scRNA-seq markers of gRSC layer 4 for two replicates, in space (left and middle columns respectively) and summarized across arc-length distance from RSC (right). B-D. As in (A), but for gRSC layer 4 marker genes (B), VIS layer 2/3 marker genes (C), and VIS layer 4 marker genes (D). Scale bar: 1 mm.

