## Supplemental Figure 1 for "Sharp cell-type-identity changes differentiate the retrosplenial cortex from the neocortex"

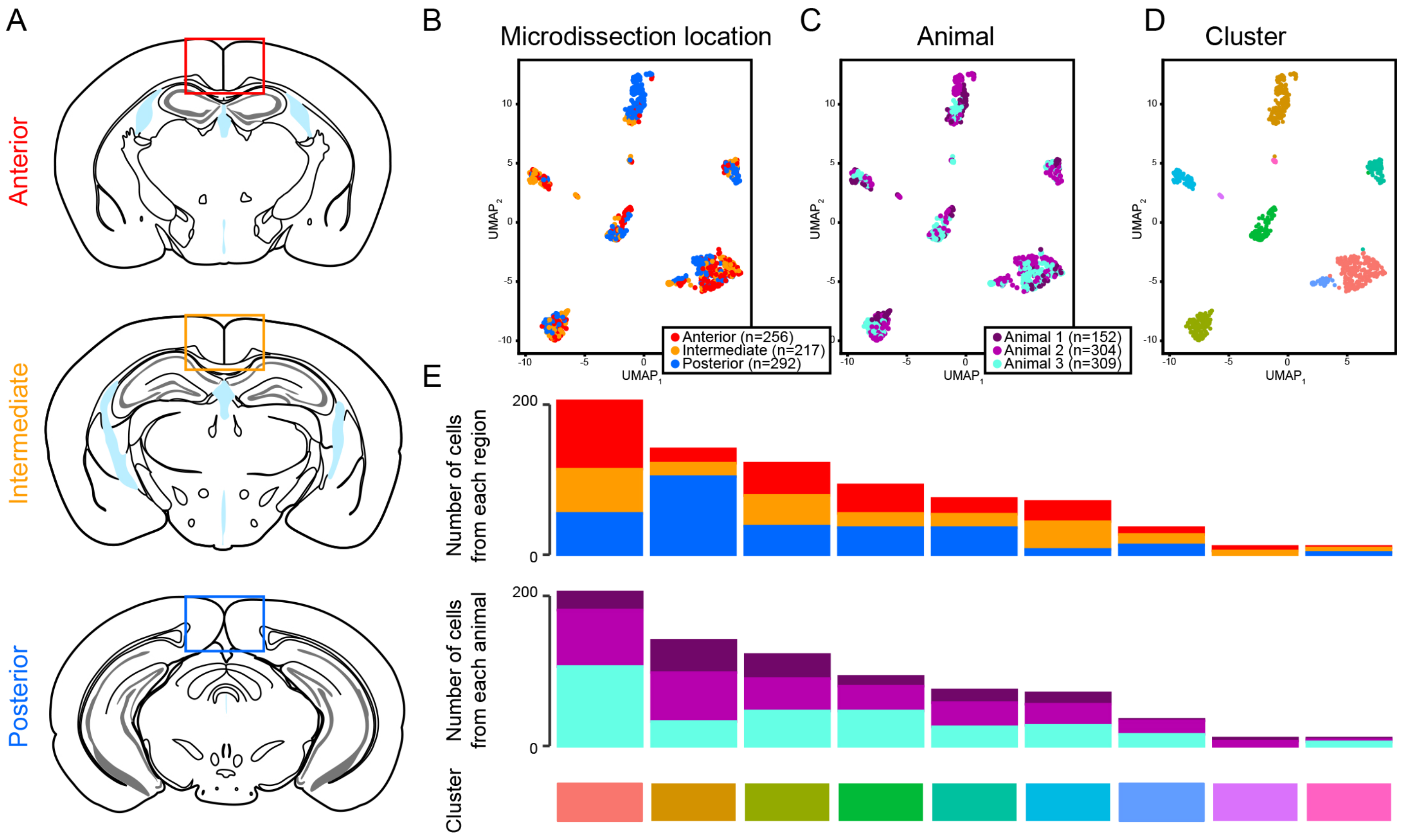


**Figure S1. Consistency of single-cell transcriptomes across spatial locations and animals.**
Related to Figure 1.
A. Schematized location of microdissected anterior, intermediate, and posterior locations. B. UMAP dimensionality reduction showing microdissected region for each cell. C. As in (B), but showing original animal for each cell. D. UMAP dimensionality reduction showing cluster identity, for comparison to (B,C). E. Quantification of number of cells for each cluster.
