## Supplemental Figure 2 for "Sharp cell-type-identity changes differentiate the retrosplenial cortex from the neocortex"

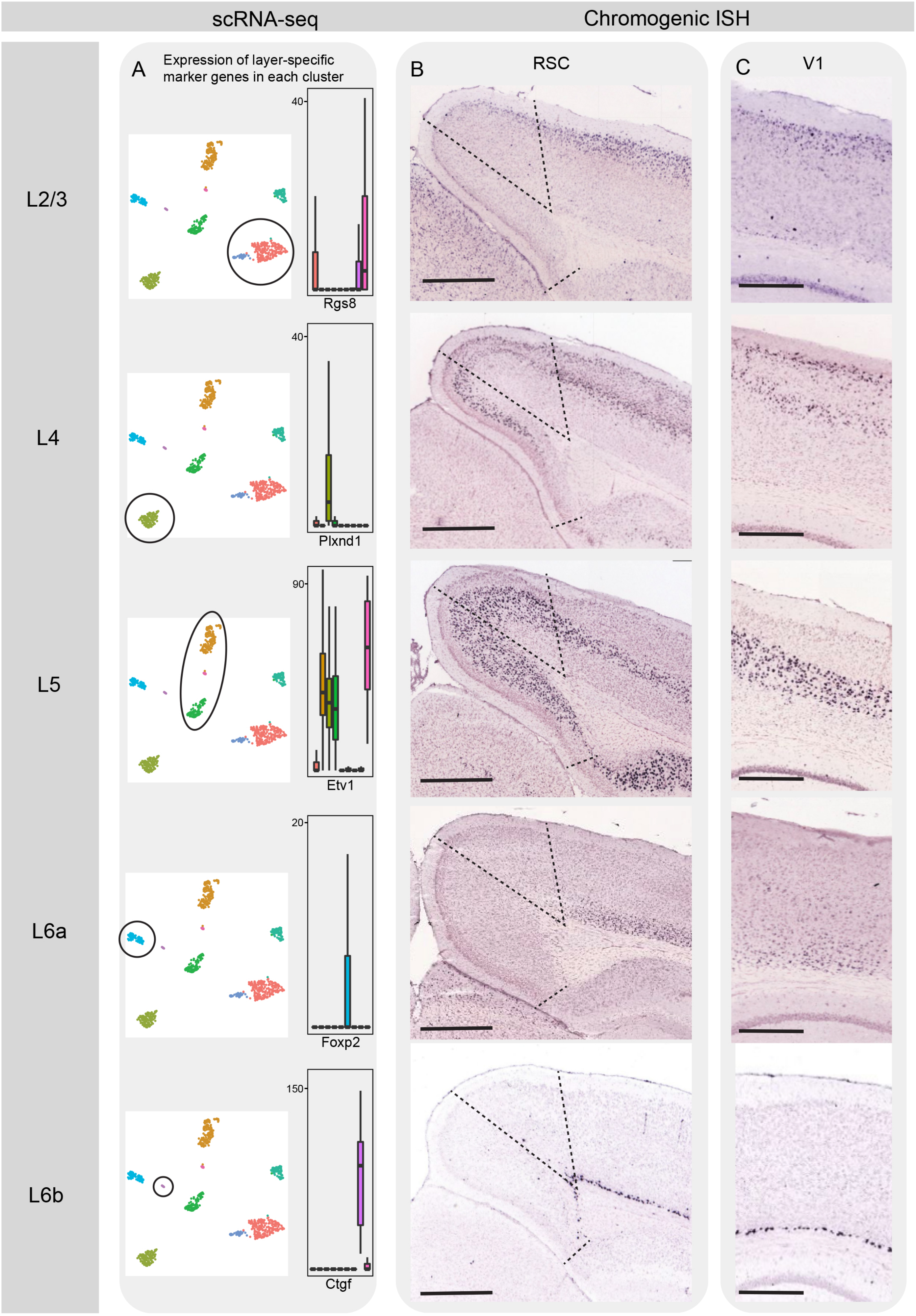


**Figure S2. Spatial mapping of layer-specific marker genes in the RSC and neocortex.**
A. Expression of laminar marker genes in scRNA-seq RSC dataset (scale bar: 839μm). B. Expression of laminar marker genes in chromogenic ISH dataset in RSC (scale bar: 419μm). C. As in (B), but for region V1 to compare layer-specific labeling to RSC. Individual rows illustrate marker genes for different cortical laminae.
