## Supplemental Figure 3 for "Sharp cell-type-identity changes differentiate the retrosplenial cortex from the neocortex"

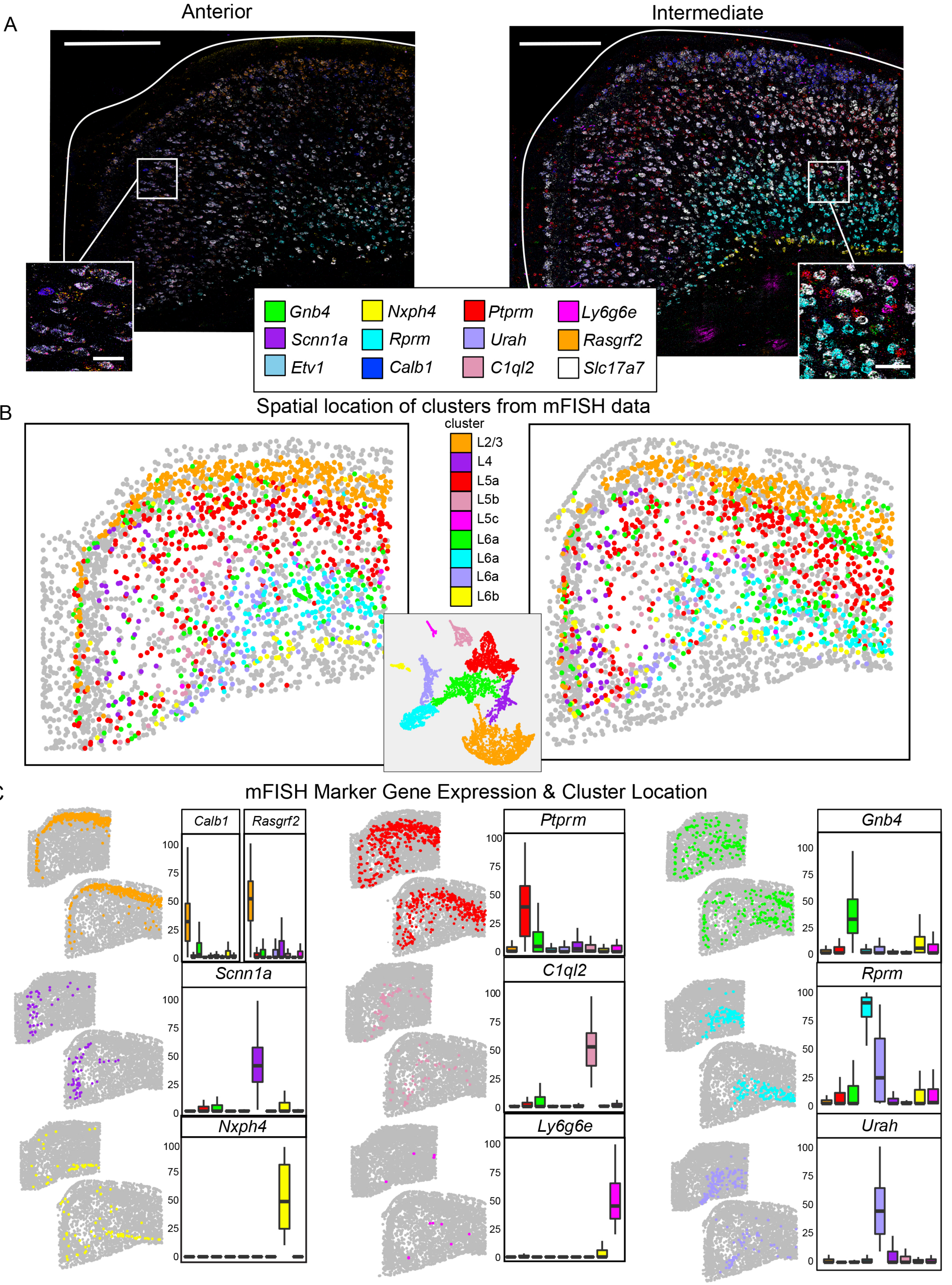


**Figure S3. mFISH clusters and marker genes across the anterior-posterior axis of the RSC.**

Related to Figure 3.

A. mFISH images of anterior and intermediate RSC (scale bar: 300µm), with insets showing expanded regions (scale bar: 50µm). Legend provides marker genes and coloring conventions. B. Clustering of mFISH data for anterior and intermediate sections. Inset provides UMAP for reference. C. Individual clusters in space, with boxplots providing marker gene expression for each cluster.
