## Supplemental Figure 4 for "Sharp cell-type-identity changes differentiate the retrosplenial cortex from the neocortex"

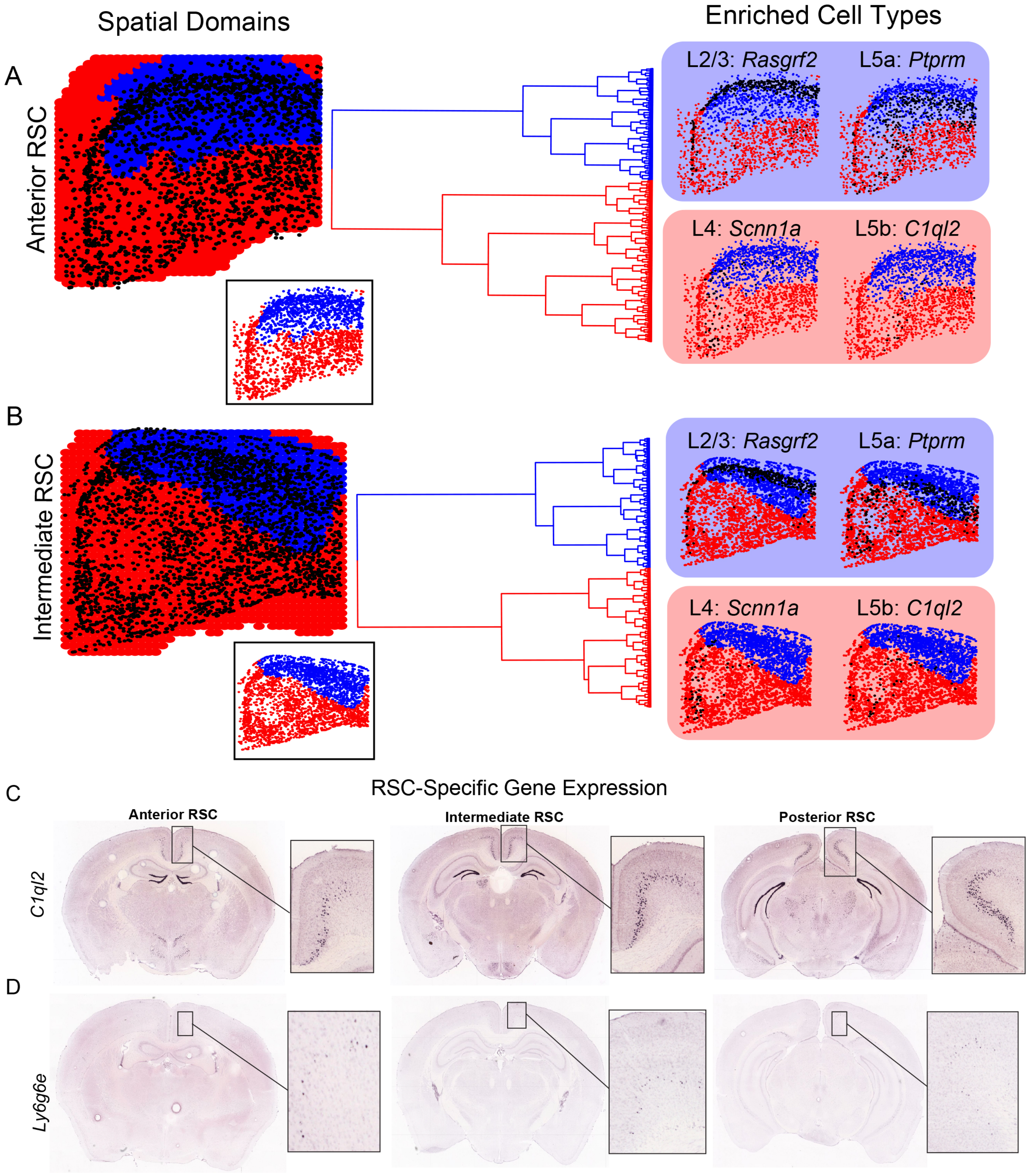


**Figure S4. Subregions across the anterior-posterior axis of the RSC.**
Related to Figure 4.
A. Spatial domain analysis of the anterior RSC, with hierarchical clustering and cell type composition analyses as in Figure 4. B. As in (A), but for the intermediate RSC. C. Expression of C1ql2, an RSC-specific marker gene of L5b, across the anterior-posterior axis of the RSC. Image from the Allen Mouse Brain Atlas. D. As in (C), but for the RSC-restricted L5c marker gene Ly6g6e.
