## Supplemental Figure 5 for "Sharp cell-type-identity changes differentiate the retrosplenial cortex from the neocortex"

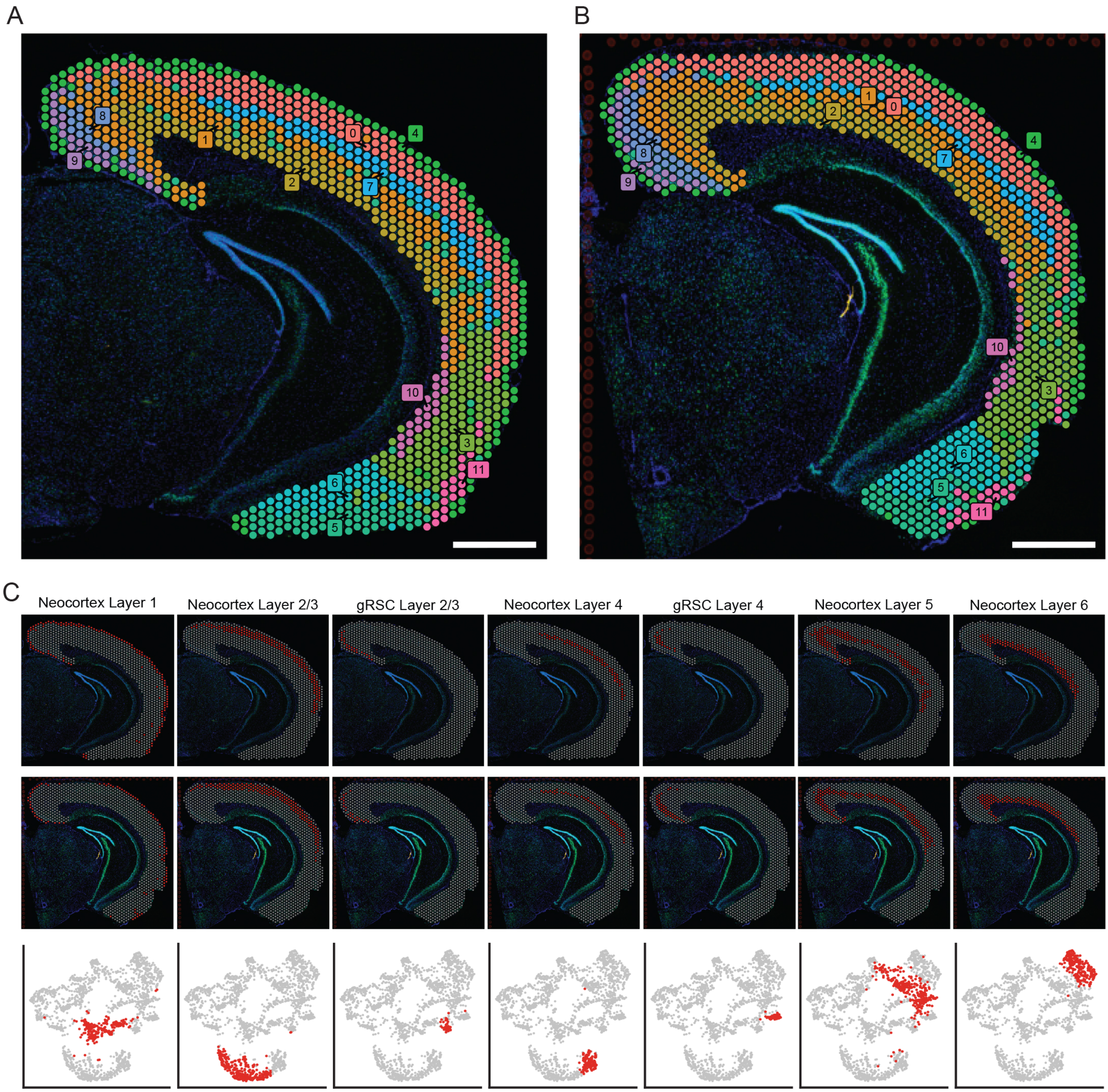


**Figure S5. Reproducibility of spatial transcriptomics clusters.**
Related to Figure 6.
A,B. Spatial transcriptomics data colored according to cluster identity for 2 replicates (A and B, respectively). Scale bar: 1 mm. C. Individual clusters highlighted in red spatially shown for two replicates (top and middle rows, respectively), with UMAP dimensionality reduction provided for reference (bottom row).
