## Supplemental Figure 6 for "Sharp cell-type-identity changes differentiate the retrosplenial cortex from the neocortex"

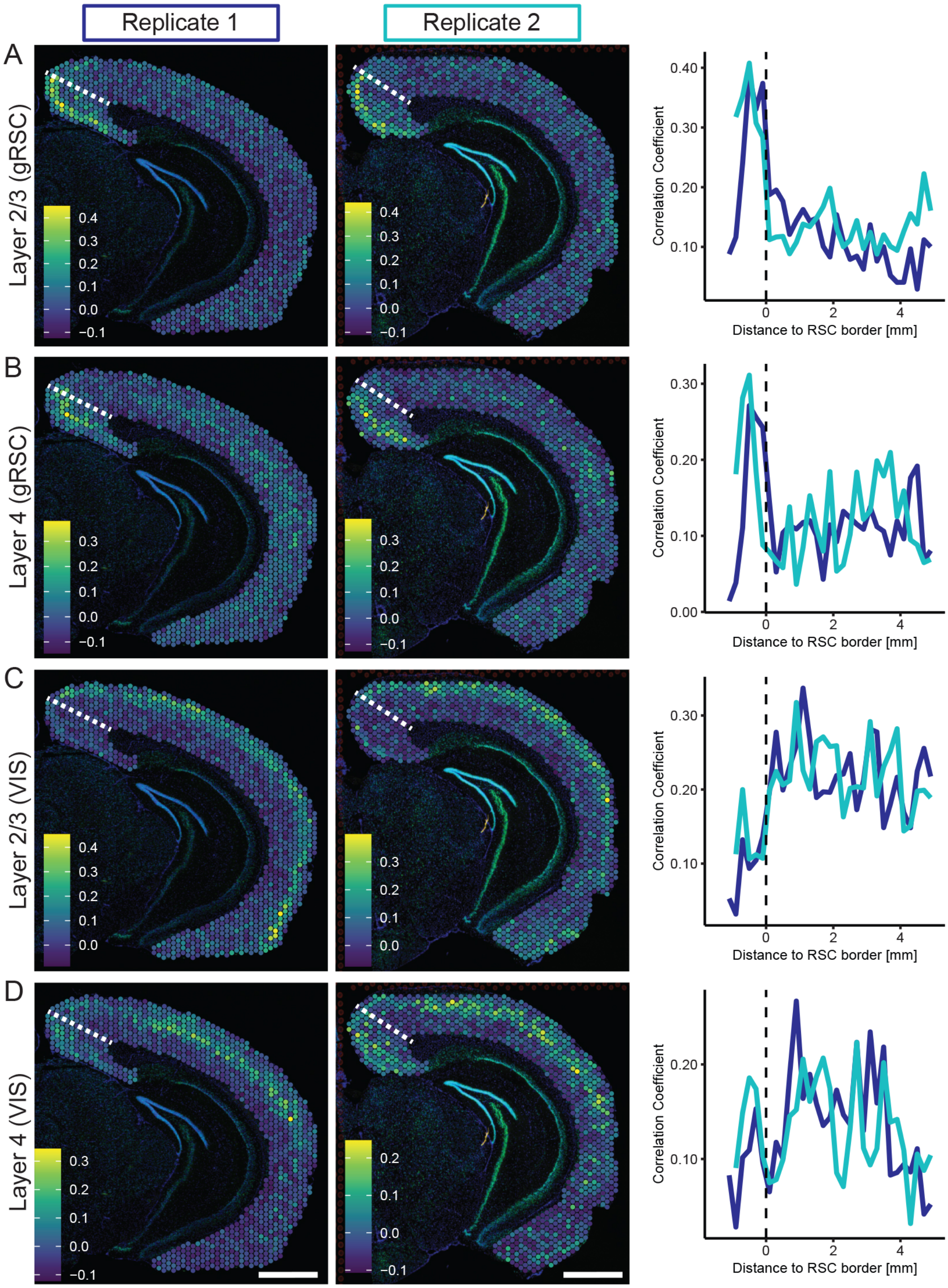


**Figure S6. Reproducibility of region-specific correlation of ST and scRNA-seq.** Related to Figure 7.

A. Pearson correlation coefficient between ST data and average gene expression of scRNA-seq markers of gRSC layer 4 for two replicates, in space (left and middle columns respectively) and summarized across arc-length distance from RSC (right). B-D. As in (A), but for gRSC layer 4 marker genes (B), VIS layer 2/3 marker genes (C), and VIS layer 4 marker genes (D). Scale bar: 1 mm.
